# Astrocytes in white matter respond to tensile cues during cortical folding: a numerical study

**DOI:** 10.1101/2025.10.17.683172

**Authors:** Karan Taneja, Kengo Saito, Hiroshi Kawasaki, Maria A. Holland

## Abstract

Our understanding of the process of formation of gyri (ridges) and sulci (furrows) in the cerebrum during development is moving beyond the role of neurons. Glial cells such as astrocytes, which are the most common cell type in the brain, are especially prominent under the gyri and have been shown to be essential for gyrification in ferrets. Their dysfunction has been linked to a host of neurodevelopmental diseases and disorders in humans, leading to abnormal folding patterns, hence, it is crucial to understand their role in the mechanics of folding. In this work, we propose two hypotheses of how astrocytes affect cortical folding. Our previous study demonstrated that astrocytes proliferate in the white matter (subcortex), and that inhibiting this process impairs gyrification. This leads to the ‘*pushing*’ hypothesis, where astrocytes push up the cortex outwards, leading to formation of folds. On the other hand, *ex vivo* studies demonstrate areas of the cortex and subcortex that experience tension, due to the differential growth between materials in the brain tissue. This leads to the ‘*pulling*’ hypothesis, where astrocytes experience tension from the surrounding tissue, leading to their proliferation, distribution of tensile stresses, and initiation of growth in those regions. Using the theory of finite growth, we implement these hypotheses via morphogenetic growth (*pushing*) and stress-driven (*pulling*) growth criteria. We find that morphological trends during development between the *pushing* and *pulling* effects are not dissimilar, with a comparable gyrification index. The stress distributions from both models also show common features, but the *pulling* effect shows tension in the subcortex, which matches trends observed in experiments. Therefore, it is more likely that the astrocytes affect gyrification by proliferating as a response to tensile cues and decrease the tension they experience, leading to deeper folds, rather than astrocytes independently proliferating under a gyri and then pushing the cortex up.

## 1 Introduction

The formation of folds in the cerebrum during development is a sophisticated phenomenon that leads to the formation of gyri (ridges) and sulci (furrows). The complexity of this process, which is called gyrification, results mainly from the bio-chemo-mechanical interactions between multiple cell populations and the surrounding substrate during development (Wang et al. 2022; Zarzor et al. 2023). Across gyrencephalic species (those with folded brains), cortical folds allow an increase in the surface area of the brain, accounting for superior information-processing capabilities compared to lissencephalic species (those with smooth brains). It has been predicted that morphological abnormalities, such as lissencephaly and polymicrogyria, arise from dysfunctions during the biological processes involved in cortical folding such as genesis, proliferation, migration, etc. (Budday et al. 2014).

While the initial focus was on the role of the skull constraining the growing brain, our understanding of gyrification has since progressed to investigate the interactions between neurons and the surrounding tissue (Garcia et al. 2018). It has been theorized that tangential growth in the cortex, caused by neurons proliferating from the ventricular zone and migrating outwards to form the cortical plate, generates stress and leads to folding (Bayly et al. 2013b; Bayly et al. 2014; Holland et al. 2017; Wang et al. 2021; Budday et al. 2014; Shinmyo et al. 2017). Gyrification could additionally be influenced by the tension generated by axonal connections (Van Essen 2020; Wang et al. 2024), local zones of increased neuronal proliferation (Reillo et al. 2011; Kriegstein et al. 2006; Juan Romero et al. 2015; Matsumoto et al. 2017; Matsumoto et al. 2020), or other phenomena.

While most research has focused on the role of neurons in gyrification, glial cells, including astrocytes, are the most numerous cell type in the brain. Their roles include providing structural support to neurons, maintaining the blood-brain barrier, and regulating synaptic functions (Tabata 2015). Crucially, glial dysfunctions have been linked to neurodevelopmental disorders such as Autism Spectrum Disorder (Kim et al. 2020; Unnisa et al. 2023; Neniskyte and Gross 2017), as well as Fragile-X and Rett syndrome (Jacobs and Doering 2010; Sloan and Barres 2014). The mechanisms behind these diseases are not well understood due to limited mechanistic understanding of these glial cells during neurodevelopment.

Specifically, there is very limited understanding of the role of glial cells in cortical folding, although evidence suggests that they are a crucial component of the process. Despite the traditional focus on neurons, they are not sufficient to produce folds by themselves (García-Moreno et al. 2012; Shinmyo et al. 2022). Furthermore, gyrification happens concurrently with astrogenesis, and only after neuronal proliferation and migration have ended(Neal et al. 2007; Kroenke and Bayly 2018; Rash et al. 2019), which suggests that other cells such as astrocytes (Shinmyo et al. 2022; Rash et al. 2019) are necessary for fold formation. Given this importance, it is critical to understand the mechanical role these cells play during folding, which in the future could lead to improved diagnostics, treatments, and prevention of folding abnormalities.

Ferrets (*Mustela putorius furo*) are a common animal model for the study of gyrification because unlike the mouse cerebrum, the ferret cerebrum has cortical folds. Furthermore, gyrification in ferrets occurs postnatally, making it more accessible to both imaging and interventions (Shinmyo et al. 2022). Additionally, the processes of ferret and human neurodevelopment are similar (Gilardi and Kalebic 2021; Imamura et al. 2024), although it happens on a shorter timescale in ferrets due to their 42-day gestational period. During prenatal development, neuronal progenitors are detected in the neocortex and start differentiating into neurons (neurogenesis). Around the same time, between embryonic day 31 (E31) and postnatal day 0/1 (P0/P1), outer radial glial (oRG) cells are also detected in the germinal zone (GZ) (Toda et al. 2016) as it expands in volume (Gilardi and Kalebic 2021; Matsumoto et al. 2020). Chemical signatures of glial cells start to be detected at P0, with astrocytes strongly detected around P6 (Gilardi and Kalebic 2021; Reillo et al. 2011). Neurons proliferate and migrate earlier than astrocytes, eventually forming the cortical plate (Wang et al. 2022; Gilardi and Kalebic 2021), with shallow folds developing around P6. Deeper cortical folds show up by P16 as astrocytes proliferate, becoming the most common cells in the brain (Gilardi and Kalebic 2021; Shinmyo et al. 2022). Gyrification mostly reaches completion by P36.

*In silico* methods such as the finite element method offer a rapid, low-cost, and non-invasive way to investigate and test hypotheses about the mechanisms of different cell types in cortical folding. These models allow for simulations of complex biological processes such as fiber reorientation, electromechanics, and tissue growth in a wide range of soft tissues (Buganza Tepole et al. 2011; Peirlinck et al. 2021) including the brain (Wang et al. 2022). Some well-established models for cortical folding have offered crucial insights regarding the influence of neuronal cell migration (Wang et al. 2022; Zarzor et al. 2024), axonal tension (Wang et al. 2024; Chavoshnejad et al. 2023), cerebrospinal fluid pressure (Jafarabadi et al. 2023), and curvature (Wang et al. 2021) on folding patterns. Simulations can be validated by comparing *in silico* morphology, axon tracts, cell densities, and stress patterns to those found in medical images(Budday et al. 2014; Solhtalab et al. 2025; Wang et al. 2022).

In this work, we propose two hypotheses for how glial cells, particularly astrocytes, affect cortical folding. The first is motivated by our recent findings that suggest that gliogenesis is crucial for gyrification. In Matsumoto et al. (2020), it was found that oRG cells accumulate under prospective gyral regions, and that suppressing the production of these cells impacts the depth of the folds formed during gyrification. Later, in Shinmyo et al. (2022), we found that locally reducing the number of astrocytes, without altering the presence of neurons, prevents the formation of gyri in ferrets. This leads to the ‘*pushing*’ hypothesis, where proliferating astrocytes push up on the cortex outwards, leading to formation of folds.

The second hypothesis is motivated by evidence that subcortical regions experience tension during cortical folding (Xu et al. 2010; Xu et al. 2009; Balouchzadeh et al. 2025). Furthermore, astrocytes are known to be mechanosensitive, suggesting that they might respond to subcortical tension (Katiyar et al. 2017; Turovsky et al. 2020). This leads to the ‘*pulling*’ hypothesis, where astrocytes experience tension from the surrounding tissue, leading to their proliferation locally; this results in volume growth and the dissipation of tensile stresses in those regions. Here, we use the theory of finite growth (Rodriguez et al. 1994) to implement *in silico* models of both of these hypotheses, and compare the resulting morphology and stress patterns to understand the role of astrocytes in cortical folding.

## 2 Methods

### 2.1 Kinematics of finite growth

We first consider the reference configuration of arbitrary material points ***X*** = [*X, Y*] within a body Ω. As the referential body Ω undergoes a motion to reach the current configuration Ωt at time *t*, the current position of the material points is defined as ***x*** = *χ*(***X***, *t*) = [*x, y*]. The mapping between the two configurations is given by a deformation gradient,

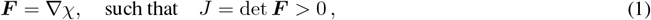

where ∇ denotes the gradient with respect to the material point ***X*** in the reference configuration.

Following Rodriguez et al. (1994), we adopt the multiplicative decomposition of the total deformation gradient to account for growth in the body,

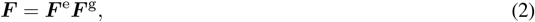

where ***F*** ^g^ is the irreversible growth part of the deformation, which maps from the reference configuration Ω to the intermediate stress-free configuration Ω_g_. Meanwhile, ***F*** ^e^ is the reversible elastic part of the deformation, mapping from the intermediate to the current configuration Ωt and ensuring compatibility. In a similar manner, the volumetric change can be decomposed into elastic and growth parts, i.e.,

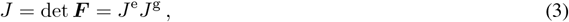

where we consider the tissue to be slightly compressible, so *J*^e^ is approximately 1.

For the specific forms of the gray and white matter growth tensors, we refer to observed phenomena in literature. Post neurogenesis in ferrets, neurons migrate to their final positions in the cortical plate (CP), causing it to grow tangentially (Reillo et al. 2011; Gilardi and Kalebic 2021). Hence, we assume that gray matter expands in the plane defined by the surface normal ***N***_**R**_,

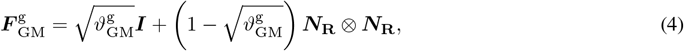

with the growth parameter representing volume change due to growth, 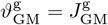.

We also assume that the subcortex expands isotropically based on previous studies (Xu et al. 2009; Bayly et al. 2013a),

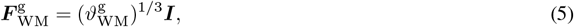

with the growth parameter that represents volume change due to growth, 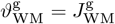

### 2.2 Constitutive equations

The cortex and the subcortex tissue are modeled as compressible neo-Hookean materials (Wang et al. 2021) with the strain energy density function

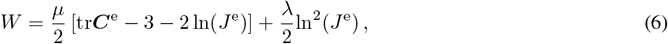

where ***C***^e^ = ***F*** ^eT^***F*** ^e^ is the elastic right Cauchy-Green tensor. The Cauchy stress tensor ***σ*** is given by

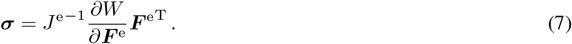

We describe the growth as quasi-static and solve the governing equation

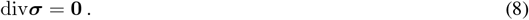

### 2.3 Growth kinetics

In this work, growth is modeled in three phases, inspired by the biological phenomena that occur during ferret development. Phase 1 corresponds to the end of prenatal development, E40-P0, where the overall volume of the brain increases (Neal et al. 2007), concurrently with the expansion of the germinal zone caused by locally proliferating glial progenitors (Matsumoto et al. 2020; Gilardi and Kalebic 2021). Furthermore, progenitors in the germinal zone are especially abundant under regions of prospective gyri (Matsumoto et al. 2020). Thus we assume that the cortex does not grow in Phase 1, but that deeper subcortical regions grow in both hypotheses, and that this growth is preferential under future gyri. To account for this spatial heterogeneity in volume expansion due to preferential presence of progenitors under future gyri, we design a spatially varying growth rate function based on two spatial parameters (Figure 2).

**Figure 1.**
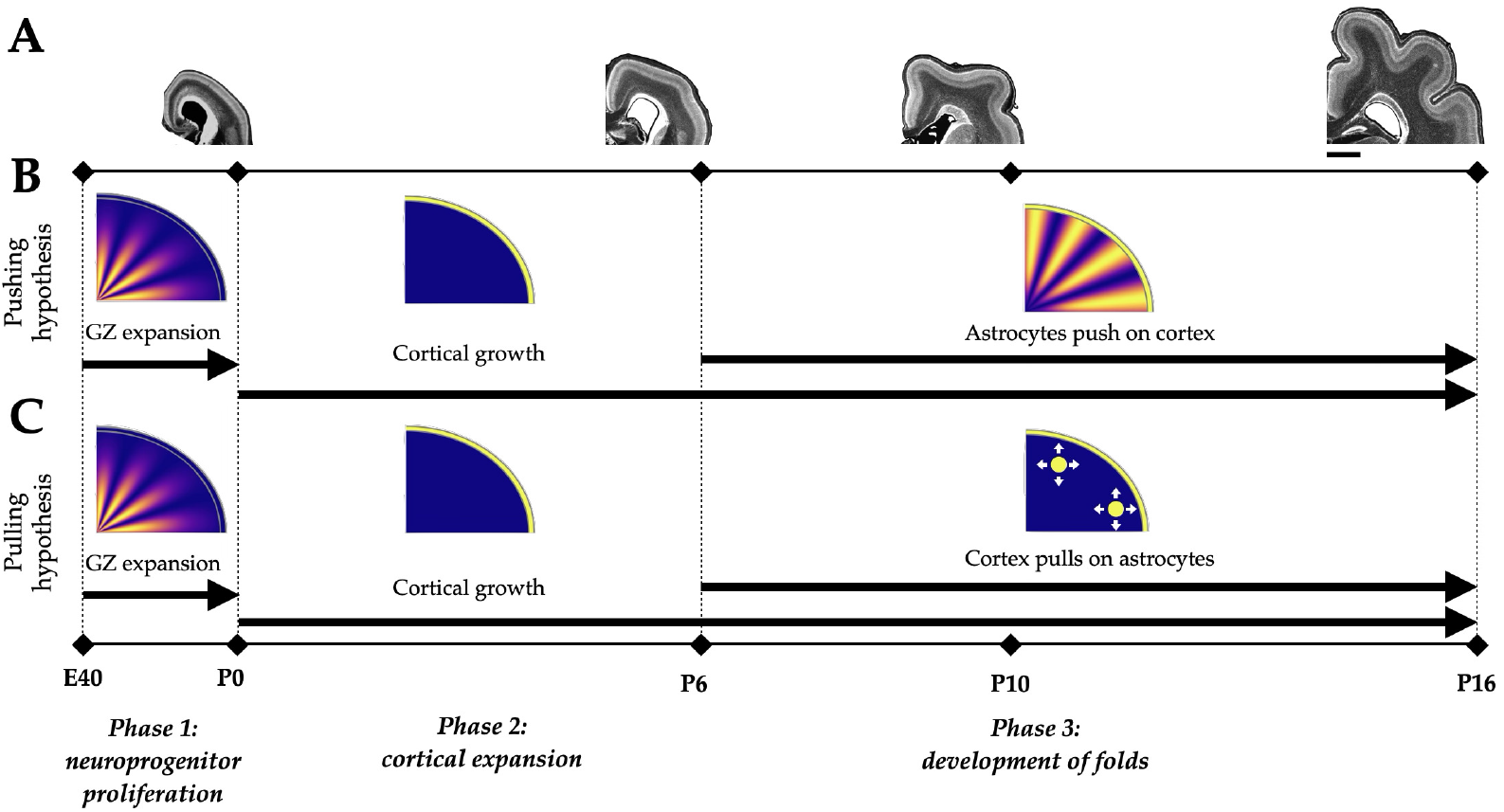
Phase definition in pushing and pulling model hypotheses. Timeline of cortical folding in ferrets used to design the phases in our model (A). Between E40 and P0, the deeper regions expand as the brain increases in size (germinal zone (GZ) expansion). Wrinkling in the cerebral cortex starts to appear around P6, with folds becoming visible around P10, and their basic structures are seen by P16. Timeline for the phases of growth in the two hypotheses in the simulation (B,C). Phase 1 models the growth of the GZ in the white matter (WM) with a larger growth rate under the prospective gyral regions. Gray matter (GM) growth is initiated in Phase 2 and continues on to Phase 3, where white matter grows according to the *pushing/pulling* hypotheses. The scale bar in (A) denotes 2 mm.

**Figure 2.**
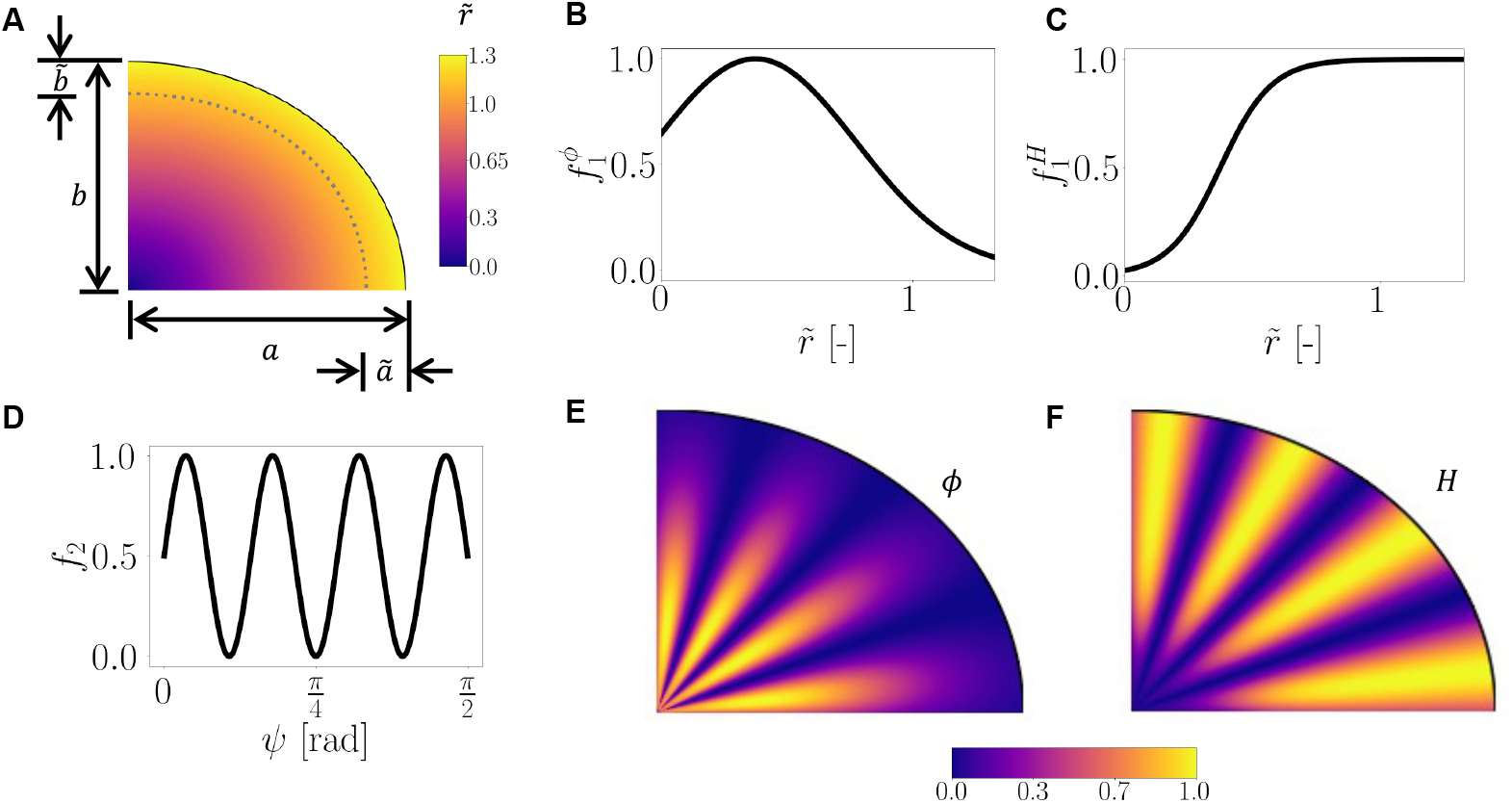
Schematics of the growth rate functions. The scaled radial distance 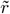 (Equation 10) (A) where the dotted line indicates 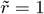. The components of the growth rate function in the white matter for Equation 12 and Equation 15 for the parameters in Table 1 (B,C and D). The resultant growth rate functions *ϕ* and *H* (E,F).

First, the angle, in radians, that a point forms with respect to the major axis is defined as

**Table 1.** Model parameters.

| Parameter | Description | Type | Value | Source/Comments |
| --- | --- | --- | --- | --- |
| $a$ | Major axis of WM ellipse | Geometric | 3.45 mm | Estimated from images |
| $b$ | Minor axis of WM ellipse | Geometric | 2.85 mm | Estimated from images |
| $t$ | Thickness of GM layer | Geometric | 0.15 mm | Estimated from images |
| $\mu_{\text{GM}}$ | GM shear modulus | Material | $2 \times 10^{-3}$ N/mm<sup>2</sup> | Wang et al. (2024) |
| $\mu_{\text{WM}}$ | WM shear modulus | Material | $1 \times 10^{-3}$ N/mm<sup>2</sup> | Wang et al. (2021) |
| $\lambda_{\text{GM}}$ | GM bulk modulus | Material | $2 \times 10^{-1}$ N/mm<sup>2</sup> | Wang et al. (2021) |
| $\lambda_{\text{WM}}$ | WM bulk modulus | Material | $1 \times 10^{-1}$ N/mm<sup>2</sup> | Wang et al. (2021) |
| $\tilde{a}, \tilde{b}$ | Reduction factors | Scaled distance function $\tilde{r}$ | 0.48 mm, 0.4 mm | Sensitivity study |
| $\alpha$ | Standard deviation | Growth rate function $\phi$ | 0.4 | Sensitivity study |
| $\delta$ | Scaled mean | Growth rate function $\phi$ | 0.415 | Sensitivity study |
| $\bar{\alpha}$ | Smoothing | Growth rate function $H$ | 10.0 | Sensitivity study |
| $\bar{\delta}$ | Scaled threshold | Growth rate function $H$ | 0.415 | Sensitivity study |
| $G_{\text{GM}}$ | Rate factor | Growth | 0.084/d | Wang et al. (2021) |
| $N$ | Number of proliferation zones | Growth | 4 | Neal et al. (2007) |

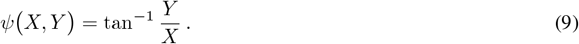

Secondly we consider 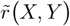, a scaled euclidean distance from the origin to the reference coordinates,

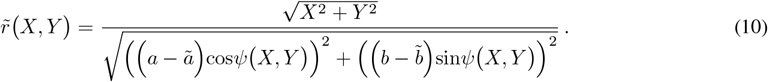

where *a* and *b* are the major and minor axes of the white matter ellipse, and *ã* and 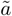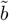 adjust the scale such that 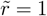at some distance inside the white matter (Figure 2A).

Thus the growth parameter 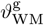 evolves as

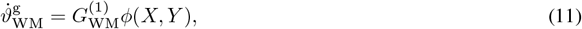

Where 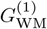is a growth rate and *ϕ*(*X, Y*) varies spatially as

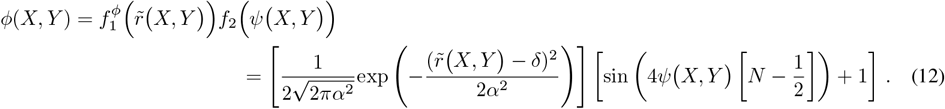

Here, the parameters *δ*and *α* indicate the mean and standard deviation of the gaussian function 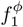, respectively (Figure 2B), and the parameter *N* denotes the number of proliferation zones, where gyri are expected to form (Figure 2D). This maintains a consistent higher growth rate in regions of higher proliferation in the deeper subcortex, with a lower growth rate near the gray-white matter interface (Figure 2E). In the absence of data on progenitor proliferation rates for the expanding subcortex in Phase 1, the growth rate of white matter in this phase is assumed to be the same as the gray matter growth rate in Phase 2, which was obtained from literature (Wang et al. 2021), i.e., 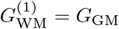.

Phase 2 corresponds to the beginning of postnatal development, P0-P6, as the cortex begins to expand. Although neurogenesis begins prenatally in ferrets, a majority of the neurons do not reach their final positions in the cortical plate until the first two postnatal weeks (Gilardi and Kalebic 2021; Wang et al. 2022). Hence we assume that gray matter starts to grow in Phase 2 of both hypotheses and continues through the end of Phase 3 (Figure 1), with a constant growth rate,

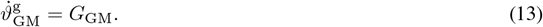

In this phase, we assume that the white matter does not expand. Although gliogenesis is detected at P0, the astrocytes that are assumed to drive growth in our model are not detected strongly until P6 in the subcortex (Gilardi and Kalebic 2021; Reillo et al. 2011), corresponding to the start of Phase 3 in our simulations (explained below).

Phase 3 corresponds to P6-P16, when folds initiate and deepen, alongside an increase in astrocyte proliferation and an overall expansion of the volume of the subcortex (Neal et al. 2007; Jacobsen and Miller 2003). Here, we assume that cortical growth continues while both the *pushing* and *pulling* hypotheses cause isotropic growth (Equation 5) in the subcortex, although the locations and mechanisms of this growth differ.

For the *pushing* hypothesis, white matter grows due to astrocyte proliferation in a similarly periodic pattern as in Phase 1,

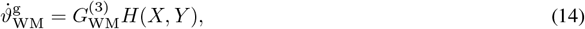

Where 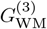is the growth rate parameter and *H*(*X, Y*) is another spatially varying growth rate function that accounts for the pushing effect of astrocytes under the gyral locations (Figure 2B),

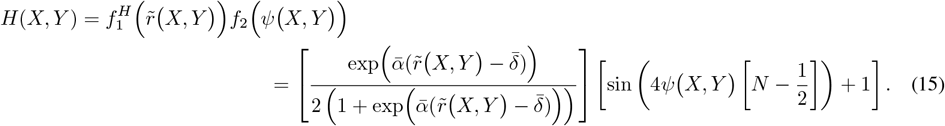

Here, the parameters 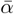, 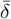are smoothening and threshold parameters in the sigmoid function (Figure 2C). The scaled radial distance 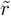maintains a higher growth rate near the gray-white interface (Figure 2F). As there are no data on astrocyte proliferation rates in gyrencephalic folding, we assume that the maximum rate of growth in the white matter in the *pushing* hypothesis is proportional to the cortical growth rate, scaled by a factor *γ*,

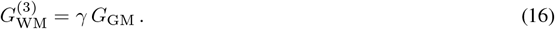

Experiments have shown that areas under growing gyri are under tension (Xu et al. 2010), which coincides with the astrocyte population increasing in the white matter (Shinmyo et al. 2022), concurrent with gray matter expansion and cortical folding. Thus, for the *pulling* hypothesis, we assume that tensile stresses initiate astrocyte proliferation, resulting in isotropic growth (Rodriguez et al. 1994; Göktepe et al. 2010; Rausch et al. 2011),

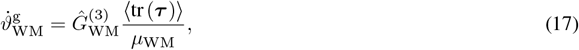

where 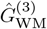 is the growth rate parameter, ⟨·⟩ is the Macauley bracket, ***τ*** is the Kirchhoff stress ***τ*** = *J****σ***, and *μ*_WM_ is the white matter shear modulus, used to normalize the Kirchhoff stress. Note that some formulations use the trace of the Mandel stress (Göktepe et al. 2010) instead of the Kirchhoff stress (Rausch et al. 2011), but they can be shown to be equivalent (Lamm et al. 2022). Here again, as the rate at which astrocyte proliferation may respond to tensile stresses in the cerebrum is unknown, we assume proportionality between the cortical growth rate and the stress-driven white matter growth rate, such that

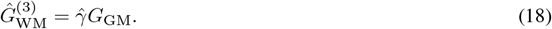

### 2.4 Numerical model

The three-phase growth model was set up in Abaqus/Explicit (Dassault Systèmes Simulia Corp. 2022) with user-defined subroutines for the two hypotheses, with the material and geometric properties given in Table 1. The simulation domain consists of a quarter of an ellipse with major axis *a* and minor axis *b* for the white matter (Figure 3). The gray matter is a strip of thickness *t* which is bound between the ellipse formed by the white matter and an ellipse with major axis as *a* + *t* and minor axis as *b* + *t*. The entire domain is meshed with ~ 18k C3D8R elements. The mass scaling factor setting in Abaqus/Explicit was selected between a range of 120-180 to allow for economical simulation times. Symmetrical boundary conditions are applied. The model geometry has a unit thickness in the out-of-plane direction with appropriate boundary conditions to mimic plane-strain conditions. To maintain the quasi-static equilibrium, the kinetic energy of the system needs to be less than 10% of the total energy, hence a small viscous pressure, 1 *×* 10^*−*5^ MPa was applied at the top surface of the gray matter. The codebase for this project is available online at https://github.com/mholla/astrocytes_pushing_pulling.

**Figure 3.**
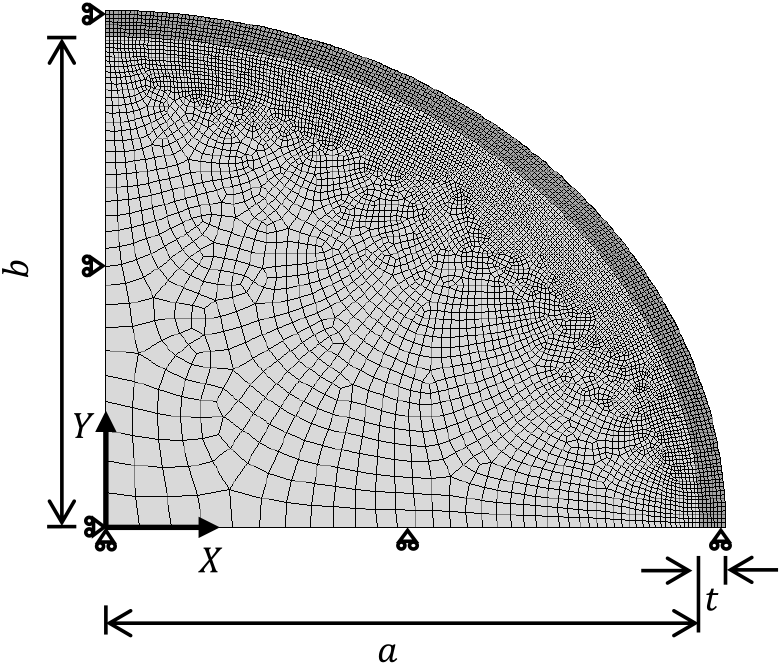
The numerical model with the finite element mesh for gray and white matter (darker and lighter shades of gray, respectively), and boundary conditions is shown for a quarter ellipse with unit thickness.

After solving, stresses can be rotated to align with radial and tangential directions of the ellipse at any point in the white matter for comparison with *ex vivo* results (Xu et al. 2009; Xu et al. 2010). At any point (*X, Y*) in the white matter, the normal vector ***N***_**R**_ can be found by taking the gradient of the ellipse equation, and its angle obtained as

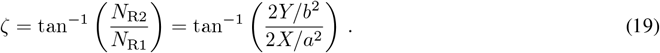

The tangential direction is found from the vector orthogonal to ***N***_**R**_. Finally, a rotated tensor 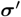 with components aligned with the local radial (normal) and tangential directions can be obtained from the stress tensor ***σ*** in the global coordinate system by

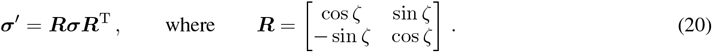

All simulations run from E40 to P16. The phases are defined based on the timeline of ferret development (Figure 1). Phase 1, in which glial progenitors proliferate locally in the expanding germinal zone, runs from E40 to P0 (approximately 10% of the total simulation time). The cortex expands tangentially after neurons reach the cortical plate in the first two postnatal weeks (Gilardi and Kalebic 2021; Wang et al. 2022) and hence, Phase 2, in which the cortex begins to grow, runs from P0 to P6 (approximately 35% of the total simulation time). Finally, while glial progenitors are present shortly after birth (Gilardi and Kalebic 2021; Jacobsen and Miller 2003), the astrocyte marker GFAP is strongly visible only after P6 (Reillo et al. 2011); therefore, it is assumed that the subcortex does not grow in Phase 2 in our model. However, in Phase 3 that runs from P6 to P16, both the subcortex and the cortex grow and the astrocyte hypotheses differentiate, with the *pushing* and *pulling* mechanisms affecting approximately 55% of the total simulation time.

### 2.5 Gyrification index calculations

To compare the morphology between simulations and experiments quantitatively, we calculated the gyrification index (GI).

From the simulations, the deformed coordinates of the outer (pial) surface were extracted, and a convex hull was created around them using the SciPy package from Python. The length of both surfaces was calculated, and the GI was calculated as

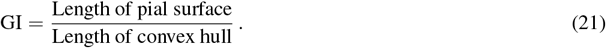

Coronal images with Hoechst stains from a ferret at P10 (*n* = 2) and P16 (*n* = 3) were post-processed with ImageJ (Schneider et al. 2012) using the thresholding procedure before extracting the coordinates of the edge of the cortical plate using the wand tool. Cubic B-splines were used to smooth the raw data, and the pial surface was obtained by manually selecting the endpoints. Then, a convex hull was fit around the pial surface, from which the GI metric was calculated as described previously.

## 3 Results

The three-phase growth model was set up as described in subsection 2.4. The white matter expands as astrogenesis and gyrification occur (Shinmyo et al. 2022; Gilardi and Kalebic 2021), but to the best of our knowledge, no studies have been conducted on the proliferation rates of astrocytes in the growing subcortex in gyrencephalic brains. Hence, we consider a range of values for growth ratios *γ*, 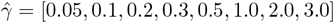, including rates of white matter growth that are both slower and faster than gray matter. Note that we consider the same range although the mechanisms for pushing (morphogenetic growth) and pulling (tensile stress-driven growth) effects are different.

### 3.1 Development of in silico folding morphology over time

We begin by considering two control cases, in order to see the effect of the two hypotheses (Figure 5). In the first, we model the growing cortex on a non-growing substrate, such that the morphology is driven purely by the buckling of the bilayer system from growth in the cortex (Figure 5A).

**Figure 4.**
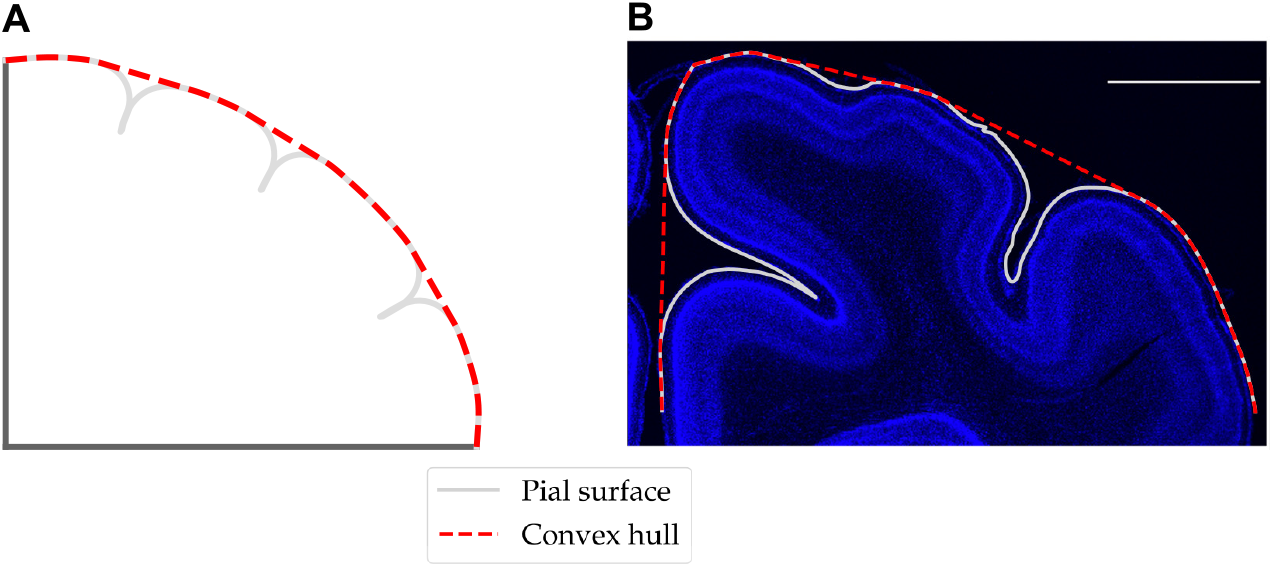
Schematic for calculation of GI using Equation 21 from simulation (A) and histological images (B). Both images are shown at P16, and the scale bar in (B) denotes 2 mm.

**Figure 5.**
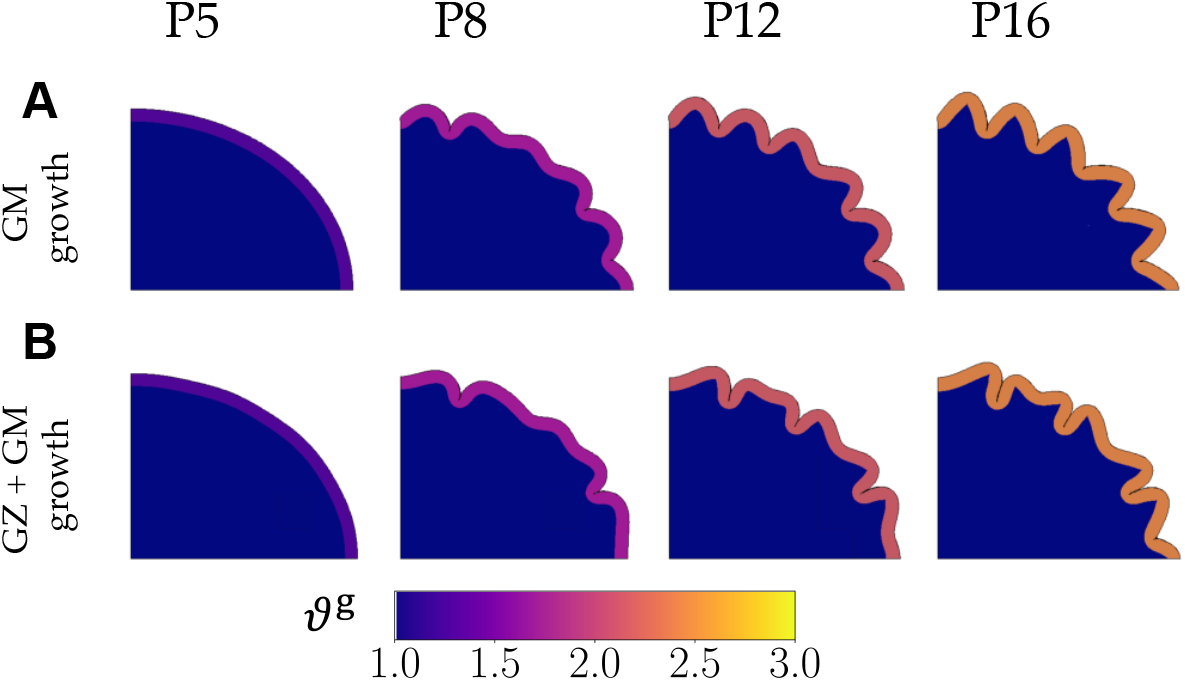
Development of folds in control simulations with neither pushing nor pulling effect. In the first control (A), only the gray matter grows (no white matter growth), and in the second control (B), germinal zone (GZ) expansion is followed by the growth in the gray matter only. The final shape in the first control case is only a result of the buckling of the bilayer system, while the shape in the second case is affected by the perturbations from growth in Phase 1.

The second control case includes GZ expansion in Phase 1 along with the growing cortex (Figure 5B). Here, white matter growth in Phase 1 happens primarily in *N* zones of proliferation, creating perturbations that modify the natural buckling pattern of the bilayer system.

In the two models of the pushing and pulling hypotheses, the zones of growth in the subcortex in Phase 1 again act as perturbations that affect the folds that result later. After the onset of Phase 3, the *pushing* effect continues to grow in the same zones as those that expanded in Phase 1, forcing the gyri to appear in those locations (Figure 6).

**Figure 6.**
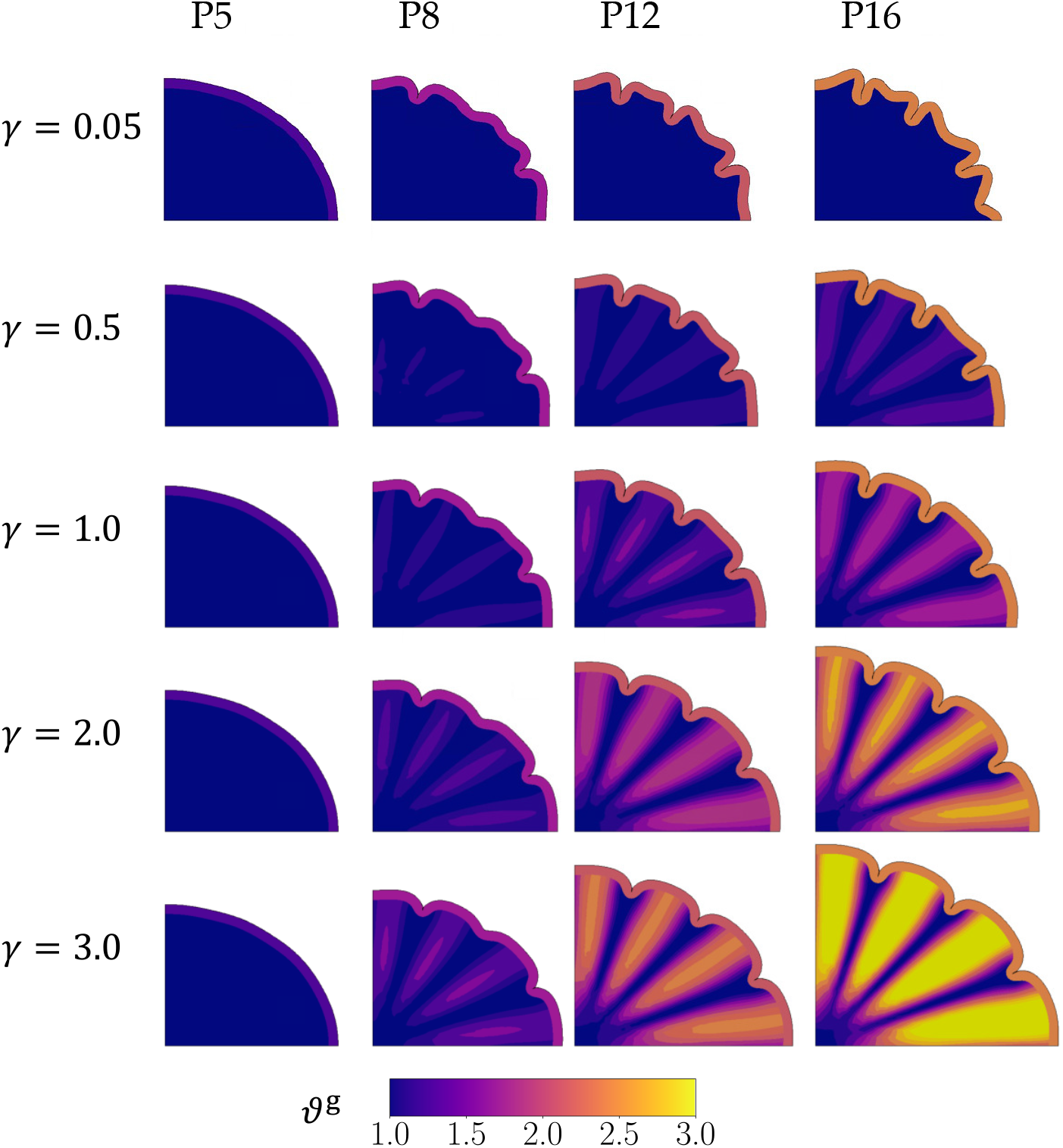
Development of folds in the *pushing* hypothesis. The first column shows the simulation in Phase 2, while the remaining are in Phase 3 of the simulation. The number of gyri matches the number of proliferation zones (*N* = 4) in all cases of the growth rate parameter *γ* considered except *γ* = 0.05 which resembles the second control case.

Similarly, the zones of growth in Phase 1 also create instabilities in the *pulling* effect, influencing the formation of gyri as Phase 3 progresses (Figure 7). The gyral locations in the *pulling* effect match the locations of progenitor proliferation for a significant range of the white-gray growth ratio parameter (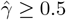).

**Figure 7.**
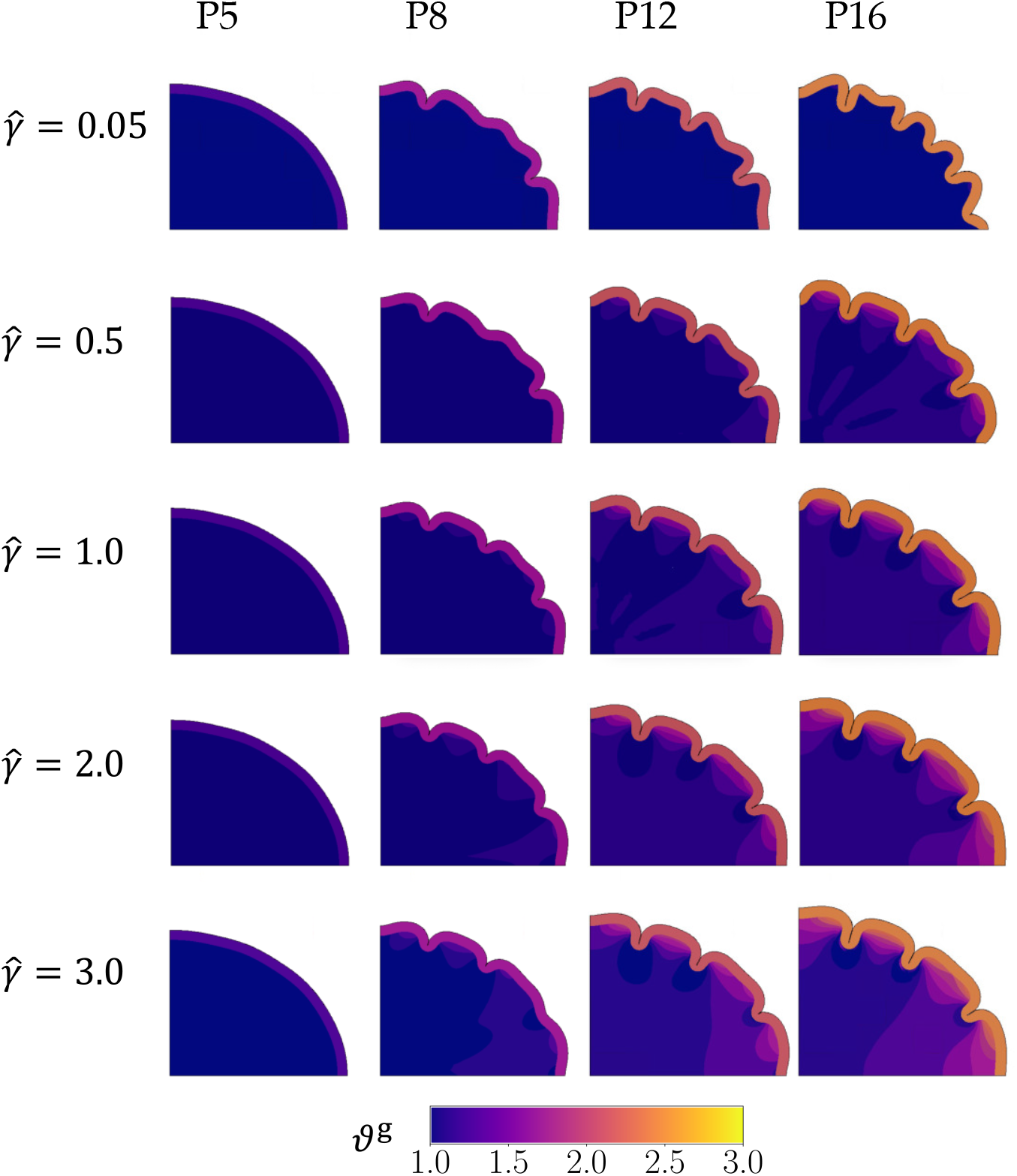
The development of folds for the parameters in the *pulling* hypothesis are shown. The first column shows the simulation in Phase 2, while the remaining are in Phase 3 of the simulation. The number of gyri matches the number of zones of proliferation (*N* = 4) for an optimal range of the growth rate parameter, i.e., for 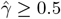.

To further quantify the morphological features, we plot the gyrification index (GI) for all growth ratios from the two models, and the control cases throughout development (Figure 8). They are also compared with GI values from histological images at P10 and P16 obtained as described in subsection 2.5.

**Figure 8.**
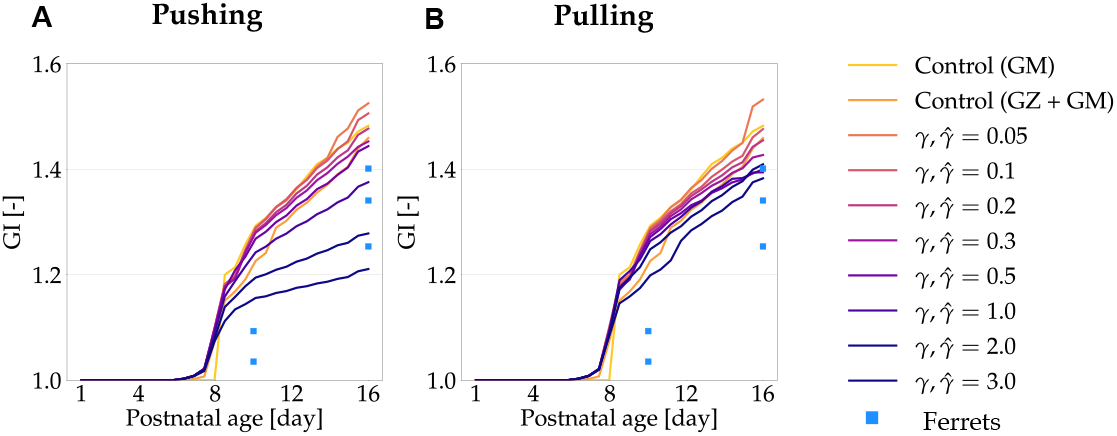
Simulated gyrification index (GI) over time for models with *pushing* (A) and *pulling* (B) hypotheses, compared to metrics from control cases and histological images. The data from P10 and P16 are taken from histological images of ferrets and are represented as square markers. *In silico* results from both hypotheses show that morphological changes happen around P6, with the GI results observed to be within the range of histological data at P16 for some growth rates.

### 3.2 Stress distributions for push and pull effects

To further compare the two models, we obtain stresses in radial and tangential directions at each point in the white matter from the simulation by rotating the stress tensor at P16 (subsection 2.4). These stresses, scaled by the shear modulus of the subcortex (Figure 9), are compared with *ex vivo* studies (Xu et al. 2010; Balouchzadeh et al. 2025) and discussed in the next section.

**Figure 9.**
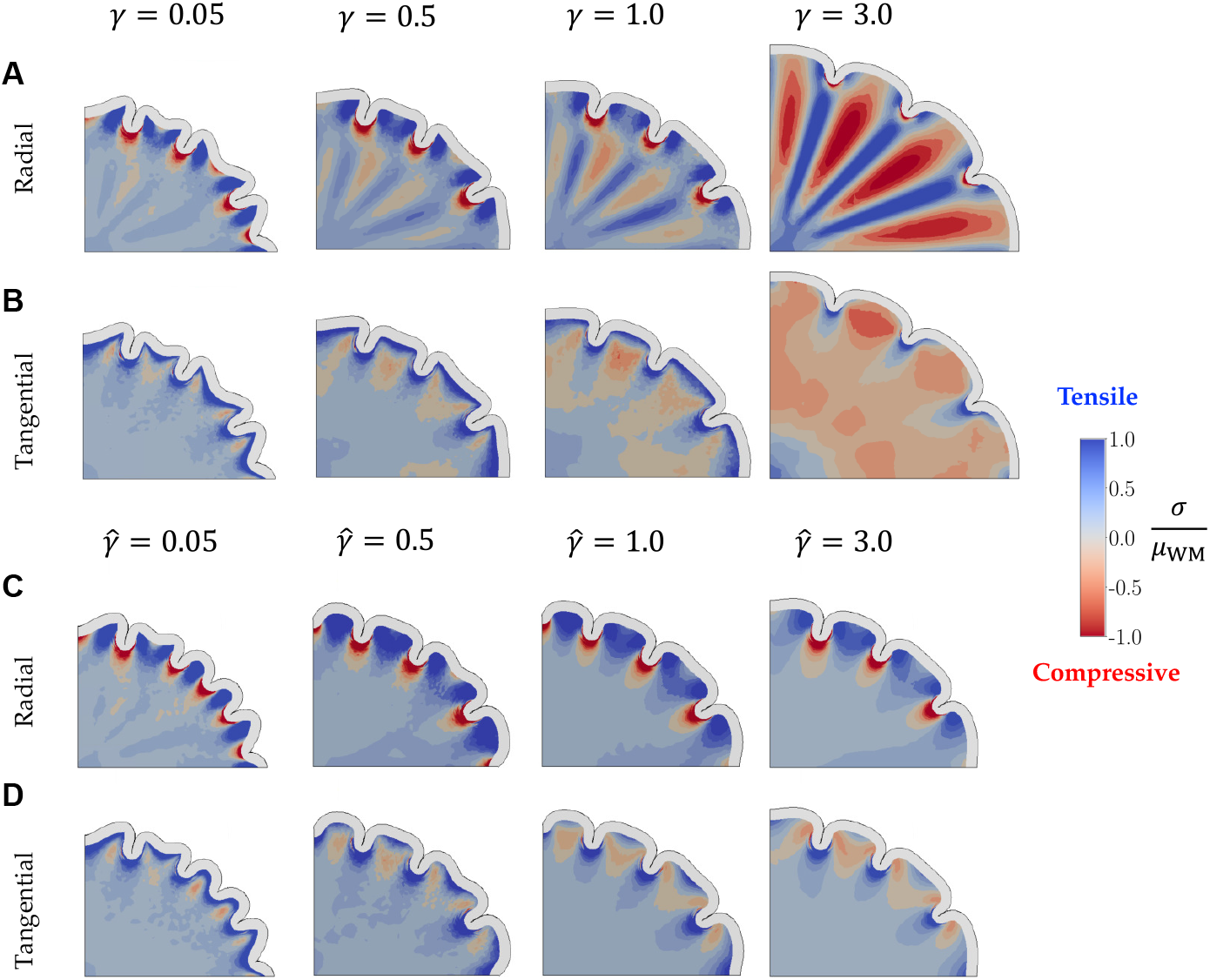
Stress distributions of the white matter in simulations of cortical folding. Final morphology (at P16) is shown along with the distribution of radial and tangential stresses for models incorporating pushing (A, B) and pulling (C, D) hypotheses. While some characteristics of *ex vivo* results are observed in both hypotheses, tangential compression observed for the pushing hypothesis in the deeper subcortex is contrary to experimental evidence.

## 4 Discussion

### 4.1 Initial perturbations from Phase 1 affect future folding patterns

Phase 1 was introduced to approximate the volumetric expansion of the GZ region during development, with higher density of oRG progenitor cells under regions of future gyri (Matsumoto et al. 2020). Since this phase is brief (only 10% of the total time), the growth caused by it is small (zones of growth show a maximum of *ϑ*^g^ = 1.15). However, these small, heterogeneous spots of growth serve as perturbations to initialize instabilities from within the deeper subcortex. The effect of these zones of positive growth in the subcortex is especially visible in the differences between the folding patterns of the control cases. For the push and pull effects, this heterogeneous growth acts as a perturbation for the respective mechanisms to act upon, guiding the final buckling pattern.

### 4.2 Folding dynamics of push and pull effects during Phase 3 progression

Following Phase 1, both models experience growth in the cortex as the gray matter expands. As the two hypotheses are not differentiated until Phase 3, both models show identical morphologies up until the end of Phase 2. For all simulations with the *pushing* effect in Phase 3, shallow folds form near the major and minor axes where two of the four zones of proliferation exist (shown at P8 in Figure 6). The models also develop smaller instabilities above the remaining two zones of proliferation. By P12, these shallow folds and instabilities progress into distinct folds, resulting in the expected number of gyri for a majority of the range of growth rate scaling parameter γ. At rates where white matter proliferation zones grow much slower than the gray matter (γ = 0.05), white matter growth is almost negligible, and the buckling cortex remains unaffected by the heterogeneous growth. This leads to a shape at P12 and P16 that is similar to the second control case, which has no pushing effect in Phase 3.

In simulations with the *pulling* effect, the cortex buckles occurs near the major/minor axes after Phase 3 begins (shown at P8 in Figure 7). However, the overall morphology shows subtle differences depending on the white matter tensile growth rate. At P8, lower growth rates of white matter (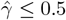) produce small buckling instabilities in the remaining cortex. At the same timepoint, an increase in the white matter growth rate (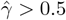) causes the instabilities to develop into folds of varying depths. By P12, similar to the *pushing* effect, simulations for all white-gray growth ratios form distinct folds. The biggest difference in shape occurs when the pulling effect is minimal (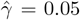), where some proto-sulci begin to emerge at P12 before fully forming at P16. Here, the natural buckling phenomenon of the system dominates over the presence of perturbations, leading to a shape that shows similarities to the control cases.

### 4.3 Number of gyri at P16 mostly equals number of proliferative regions

The shapes of the first control case (without Phase 1 growth) show that the folding morphology is solely driven by the buckling of the bilayer elliptical geometry without any perturbations (Figure 5A). In contrast, the introduction of perturbations in the second control case, with four regions of proliferation (*N* = 4) in Phase 1, changes the shape (Figure 5B). This highlights the importance of these initial perturbations in determining the final folding pattern.

For the *pushing* effect, the number and location of gyri match the number and location of proliferation zones (*N* = 4) in Phase 1 and 3 (Figure 6) for a majority of the range of growth rates *γ*. At the lowest white-gray growth ratio (*γ* = 0.05), however, the final shape is similar to the second control case (Figure 5B) as the pushing effect in the white matter is not strong enough to overcome the natural instability behavior of the buckling gray matter.

For the *pulling* effect, the number of gyri and their locations also largely match the proliferation zones (Figure 7). However, at lower white matter growth rates (i.e., when 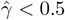), gray matter buckles faster than white matter can grow to dissipate the tension. This leads to more folds than expected at P16 resulting in a folding pattern similar to the first control case (Figure 5A).

The choice of four regions of proliferation (*N* = 4) was based on coronal histological images of ferrets from literature (Matsumoto et al. 2020; Neal et al. 2007; Barnette et al. 2009), which show four gyri. As expected, this choice of parameter results in four gyri for the majority of the simulations. To demonstrate that our model can reproduce other numbers of gyri for reasonable white matter growth rates, we tested *N* = 5 (Figure 10). Simulations with the pushing effect form five gyri for the entire range of white-gray growth parameters, while the pulling effect produces five gyri within a different optimal range (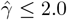). Combining this observation with the previous case (*N* = 4), there may be an upper and lower limit for tensile stress-based growth rates that results in the expected morphology.

**Figure 10.**
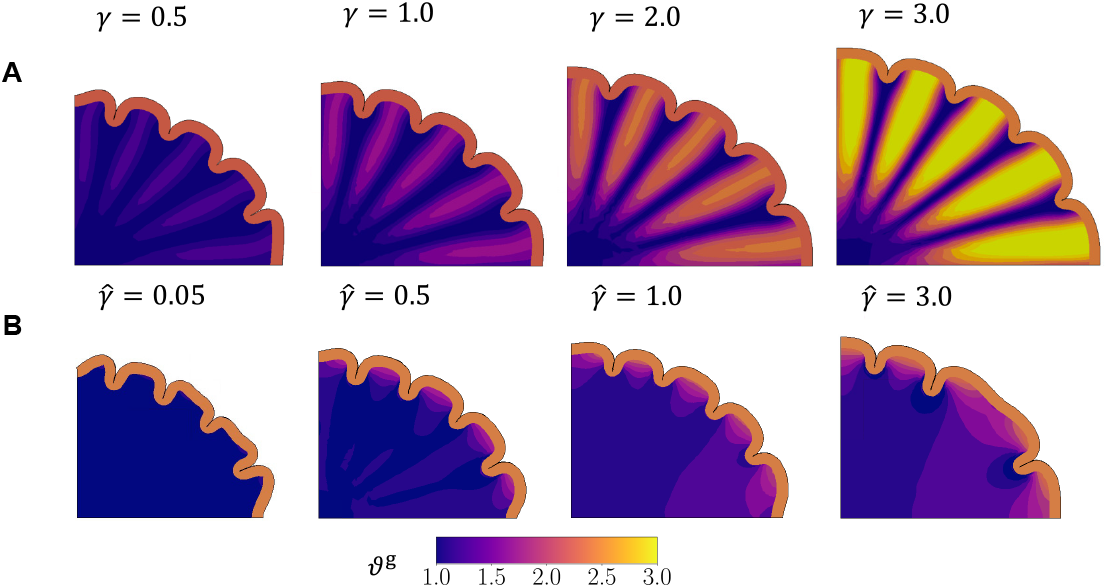
Simulated cortical morphologies during gyrification with five proliferation zones (*N* = 5) designed for *pushing* (A) and *pulling* (B) hypotheses, at the end of the simulation (P16). While all pushing effect simulations show the desired morphology, the pulling effect simulations show the expected number of gyri for a different optimal range than the case with *N* = 4, i.e., 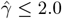.

### 4.4 The role of white matter growth rates on folding patterns at P16

Without astrocytic effects in white matter growth, the gyral shapes in the first control case (with only gray matter growth) at P16 point radially outward, driven by buckling of the gray matter layer (Figure 5A). This differs significantly from the shape of the second control case with GZ expansion followed by gray matter growth (Figure 5B). Here, growth in the white matter disrupts the natural buckling pattern of the gray matter layer, causing some sulci to form more superficially.

When white matter growth is introduced through the two models, the final shape is affected by the white-gray matter growth ratio. For the *pushing* effect, the gyri are flatter at the crowns for some slower white matter growth rates (0.5 < γ ≤ 1.0). In contrast, as white matter grows faster than gray matter (*γ >* 1.0), the gyral crowns are more rounded in shape. Such a consistent pattern does not show up in the pulling effect, with gyral crowns showing mixed curvatures across all growth ratios 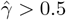.

Moreover, the expected number of gyri form for an optimal range of white matter growth rates. If the white matter growth rate is too low in both effects (e.g., *γ <* 0.5 in Figure 6 and 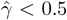 in Figure 7), it is unable to sufficiently overcome the cortical buckling effect, leading to excessive folds (similar to brains with polymicrogyria). On the other hand, if the growth rate in the pulling effect is too high (e.g., 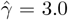 in Figure 10), it can sufficiently dissipate the building tension in the system such that buckling is impaired, and fewer folds form than expected (similar to brains with lissenscephaly). For the pushing effect, although the fastest white matter growth rate (*γ* = 3.0) leads to the expected number of gyri at P16, the degree of folding as measured by the GI at P16 in the simulation is much lower (Figure 8).

### 4.5 Gyrification index is similar between push and pull effects

Gyrification index (GI) allows us to compare our simulation results quantitatively (Figure 8). In the second control case and all of the pushing and pulling simulations, the introduction of Phase 1 growth acts as a perturbation and induces earlier buckling than the control case with only gray matter growth. Interestingly, the degree of folding as measured by GI, decreases in both models as white matter grows faster (higher *γ*, 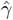). However, the variations in GI for the *pushing* effect are larger than the *pulling* effect.

From literature (Neal et al. 2007; Barnette et al. 2009), it is known that folding begins by P6, with visible folds by P10 and the basic structures fully formed by P16. Importantly, each of our simulations show that folding begins at P6, as evidenced by increasing GI. Additionally, the simulations with higher growth ratios (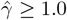) from the pulling effect and a partial range of growth ratios (*γ* = 1.0, 2.0) from the pushing effect, show similar GI values to the histological data at P16.

### 4.6 In silico stress distributions show pulling effect more likely

As morphological differences are insufficient to distinguish the two mechanisms under investigation, we obtain stresses in radial and tangential directions at each point in the white matter from the simulation, comparing the results at P16 (Figure 9).

These are compared to *ex vivo* studies (Xu et al. 2010; Balouchzadeh et al. 2025), which showed that areas under gyri should be under radial tension and those under sulci should be under radial compression and tangential tension. Further, tangential tension is also expected in the deeper subcortex as gyrification progresses (Xu et al. 2010).

In *pushing* effect simulations with slower white matter growth (e.g., 0.05 *≤ γ <* 0.5), the stress distributions closely resemble the second control case (Figure 11), where the Phase 1 perturbations only slightly influence the morphology determined by the buckling cortex. For slightly faster white matter growth rates (e.g., 0.5 ≤ γ ≤ 1.0), the gray matter effectively pulls the underlying white matter up to form folds, generating radial tensile stresses underneath the gyri. Deeper in the subcortex, however, the zones of proliferation are constrained by nearby non-growing regions, resulting in regions of compression under gyri, alternating with regions of tension under sulci (Figure 9A).

**Figure 11.**
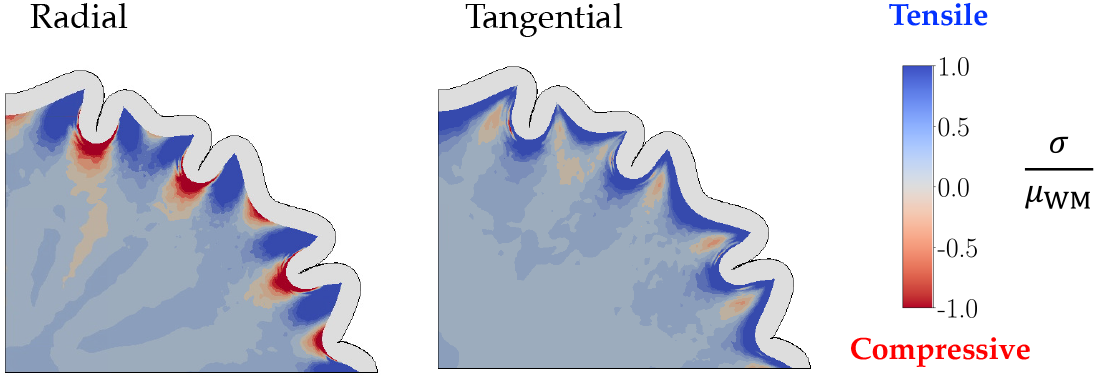
Radial and tangential stress distributions along with final morphology at P16 for the second control case are shown, where Phase 1 germinal zone expansion is followed by gray matter growth only.

As white matter growth rate increases (*γ >* 1.0), the proliferation zones in the white matter grow faster, and the cortex begins to constrain the faster-growing subcortex. This induces radial compression in the regions of growth, with tensile stresses between them where the non-growing subcortex constrains sulci. Hence, the faster the white matter grows (higher *γ*), the larger these zones of radial compression grow under the gyri. This radial compression is unrealistic.

The simulations with the *pulling* hypothesis show another consistent pattern across the entire range of growth ratios (Figure 9C). From slower tensile growth in the white matter (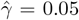), to a faster rate where the pulling effect is strong (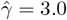), radial stresses under the gyri are tensile consistently. The tensile stresses are also larger around the crown of the gyri, and smoothly diminish to smaller values in the deeper subcortical regions.

While the radial compression under the gyri seen in the *pushing* effect at higher white-gray growth rates is unrealistic, slower white matter growth in the pushing hypothesis, as well as all cases of the *pulling* effect, show the desired radial tension under gyri. Additionally, both models show radial compression under sulci for the entire parameter ranges as well. Hence, considering just radial distributions, both effects show physiologically relevant characteristics.

We next consider the tangential stress distributions (Figure 9B,D). The differential expansion of the bilayer system creates tangential tension directly below the gray-white matter interface for a majority of the growth rates (*γ ≤* 2.0 and all 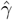), followed by a zone of compressive stresses that connect the walls of the gyrus. However, in the *pushing* effect, the zones of tangential compression continue into the deeper subcortex, contradicting the results from Xu et al. (2010), where those regions show tangential tension. Moreover, for *γ≥* 2.0, most of the domain shows tangential compression. This likely results from the isotropic growth of the white matter which, in addition to pushing radially outward, also grows to expand circumferentially and push on neighboring tissue. On the other hand, in the *pulling* effect, a majority of the subcortex is under tension in both radial and tangential directions, leading to growth in the white matter.

Considering morphology and stress distribution evidence from simulations, out of the two hypotheses, it is the *pulling* effect that matches the results from literature, and is a more likely way for astrocytes to affect cortical folding.

## 5 Conclusions

In this study, we are interested in broadly understanding the mechanics behind how glial cells such as astrocytes influence cortical folding. We developed and tested two hypotheses of astrogenesis in cortical folding, based on either a morphogenetic (*pushing*) or a tensile stress-driven (*pulling*) model of white matter growth. Simulation geometry was based on the ferret cerebrum, and our results were compared with cortical morphology and *ex vivo* stress results from ferrets during gyrification, which takes place over roughly the first two weeks postnatally.

In both the models, the number of gyri formed generally equal the number of proliferation zones, with some exceptions in the *pulling* effect outside an optimal parameter range. Overall, we find that morphological trends during gyrification between the pushing and pulling effects are not dissimilar, with comparable GI values. There are similarities in the stress distributions as well, especially in the radial direction; all pulling effect simulations, and most of the pushing effect simulations, show radial tension under gyri and radial compression under sulci, matching *ex vivo* trends (Xu et al. 2010; Balouchzadeh et al. 2025). However, the pushing effect shows tangential compression in the deeper subcortex, which is contrary to experimental evidence. Therefore, it is more likely that the astrocytes affect gyrification by proliferating in response to tensile cues, rather than astrocytes independently proliferating under gyri and pushing the cortex up.

Beyond identifying a potential mechanism for astrocyte involvement in cortical folding, our results show that the white-gray growth ratio is critical for desired folding patterns to emerge. The expected number of gyri forms within an optimal range of the tensile stress-driven growth rates in white matter, which should be sufficiently large to overcome the intrinsic buckling of the growing gray matter layer, yet low enough to not dissipate all the stress through growth. Slower rates lead to folding patterns that resemble polymicrogyria, while faster rates result in shapes that resemble lissencephaly. This aligns with evidence that dysfunctions in astrocytes lead to neurodevelopmental pathologies (Kielbinski et al. 2016; Bornemann et al. 1997), and suggests that astrocyte proliferation rates should be investigated further.

## 6 Limitations and future work

This novel study serves as a first step in understanding the role of astrocytes in the process of gyrification. It is, therefore, essential to note the limitations of this work as well. For instance, the push and pull hypotheses may not be mutually exclusive of each other. The pulling mechanism of growth attributed to astrocytes may also work in conjunction with other hypotheses of cortical folding. For example, the buckling of the growing cortex generates tensile stresses, which might affect the direction and density of white matter fiber tracts (Bayly et al. 2013b; Garcia et al. 2018; Garcia et al. 2021; Solhtalab et al. 2025), and in turn affect the placement of folds (Wang et al. 2024). Our results show that the generated tension may work alongside pre-existing patterns of astrocyte progenitors to guided astrocyte proliferation, further affecting the eventual placement of folds. The final morphology of folds maybe affected by these two working in a complementary manner, which is an area for future investigation. The generation of stresses within the subcortex may also be affected by the constraints of the skull as well as the pressure exerted by the surrounding cerebrospinal fluid (Jafarabadi et al. 2023).

As this work is preliminary, we did not consider the anisotropic and viscoelastic nature of the subcortex in our model, which could be incorporated in the future. We also focus exclusively on astrocytes, and did not consider other cells derived from oRG progenitors such as oligodendrocytes (Jovanovic et al. 2023). Furthermore, we only considered the fibrous subtype of astrocytes, and did not include the protoplasmic subtype that settles in the cortex (Tabata et al. 2022). We also do not consider any interactions between astrocytes and neurons in the cortical plate. In the future, we plan to refine this model to address these limitations.

## Acknowledgements

Research reported in this publication was supported by the National Institute of Neurological Disorders And Stroke of the National Institutes of Health under Award Number R01NS135852. The content is solely the responsibility of the authors and does not necessarily represent the official views of the National Institutes of Health. This research was also supported by commissioned research (No. 23001) from the National Institute of Information and Communications Technology (NICT), Japan; the Japan Agency for Medical Research and Development (AMED) (No. JP24wm0625112); and the Japan Society for the Promotion of Science (JSPS), KAKENHI (No. 23H00389). We also thank Dr. Johannes Krotz for their help in formulating the growth functions on an ellipse, and Julian Najera for guidance with ImageJ.

## Data availability statement

The code to run the described simulations, along with select results, are available at the online repository https://github.com/mholla/astrocytes_pushing_pulling.

